# Micromachined Thermocouple Microprobe for Real-Time Thermometry of Hippocampal Slices during High-Frequency Stimulation

**DOI:** 10.64898/2026.01.18.700190

**Authors:** Vishwendra Patel, Onnop Srivannavit, Irene C. Turnbull, Robert D. Blitzer, Angelo Gaitas

## Abstract

Neuronal activity is energetically demanding and generates heat as a metabolic byproduct. Here we investigated localized temperature changes in mouse hippocampal brain slices during intense synaptic activation. Using fast-response thermocouple microprobes placed in the tissue, we measured real-time temperature fluctuations during high-frequency (100 Hz) electrical stimulation of Schaffer collateral inputs. Stimulation induced a local temperature increase (peak ΔT = 0.54 ± 0.07 °C; n = 4). In contrast, when action potentials were blocked with tetrodotoxin (TTX), the same stimulation produced no measurable temperature rise. These results demonstrate that high-frequency neural firing and synaptic transmission can transiently elevate local brain tissue temperature, presumably due to increased metabolic activity and adenosine triphosphate (ATP) consumption. The hippocampus tolerated these small temperature shifts, but even modest heating could influence neuronal function. Our findings provide direct experimental evidence linking synaptic activity to localized brain heat production, and underscore the importance of considering thermogenesis under stimulation conditions that give rise to forms of synaptic plasticity.

## Introduction

Brain function is inextricably linked to energy metabolism and heat production^1^. Although brain temperature in mammals is often assumed to remain near a constant 37 °C, research has revealed that substantial, rapid fluctuations can occur during normal behaviors and neural activation^1–6^. Brain temperature is recognized as an important physiological parameter that reflects the metabolic activity of neural circuits and can affect neural signaling and function^1,7^. Early direct thermometry studies also documented localized temperature heterogeneity and potential gradients in brain tissue ^8,9^. For example, in awake rats, brain temperatures can vary by up to ~4 °C, dipping to ~35 °C during deep sleep and peaking near 39.5 °C during heightened activity such as exercise or copulation^1^. Increases of 1-2 °C have been documented with routine behaviors like feeding or the transition from sleep to wakefulness ^2,10,11^. These observations demonstrate that even in homeothermic animals, the brain’s thermal state is dynamic and closely tied to neural activity ^1^. Local sensory stimulation can also evoke reproducible brain temperature changes ^4,12^.

Importantly, local neural processes are a primary source of these temperature changes. During acute arousal or stimulation, brain regions often warm up before any rise in body core temperature, indicating that the heat originates within activated neural tissue rather than from peripheral sources^1,13^. This local thermogenesis is thought to result from the intense energetic demands of active circuits^1^. A key contributor is the hydrolysis of ATP required to restore ionic gradients after activity, primarily via Na^+^/K^+^-ATPases, and by other metabolic processes that collectively release heat as a byproduct^1,14^. Indeed, a substantial fraction of cellular ATP expenditure is devoted to restoring sodium and potassium gradients during neuronal signaling ^1,15^.

Heat production following electrical activity is temporally biphasic, with an initial component tightly coupled to action potentials and synaptic currents, followed by a recovery component driven by ATP-dependent restoration of ion gradients and oxidative metabolism ^14^. Consistent with classic nerve calorimetry, the initial phase can include both heat release and heat absorption, whereas the recovery phase is net heat-releasing^14,16^. At the tissue scale – particularly during high-frequency stimulation (HFS) trains – the measurable temperature transient is therefore expected to be dominated by recovery heat, because metabolic work persists beyond the duration of very brief electrical events. In addition to neuronal ion pumping, HFS places substantial metabolic demands on glial cells, especially astrocytes, which clear synaptically released glutamate through high-capacity transporters and support the glutamate-glutamine cycle; these energy-intensive processes can materially contribute to local heat generation ^17–20^. Accordingly, activity-dependent thermogenesis is best viewed as a network phenomenon that reflects coupled energetic costs across neurons and glia.

Under physiological conditions, excess heat is dissipated by cerebral blood flow; cool arterial blood acts as a thermal sink, carrying heat away and helping maintain thermal homeostasis^1,21^. This balance is often described within a bioheat framework in which tissue temperature reflects competition between intrinsic metabolic heat production and heat removal by blood perfusion ^22,23^. Consequently, in vivo measurements necessarily conflate metabolic heat generation with blood-flow-mediated cooling and neurovascular coupling. By contrast, an *in vitro* slice preparation provides a reduced model in which blood perfusion-based heat removal and neurovascular coupling are absent, enabling more direct quantification of intrinsic, activity-dependent metabolic thermogenesis than is feasible in intact animals. (We note that convective heat removal by the superfusing artificial cerebrospinal fluid persists in the brain slice preparation; however, the dominant regulated heat sink of blood perfusion and neurovascular coupling is eliminated.)

The hippocampus is a brain region of particular interest for studying activity-dependent thermogenesis. The hippocampus-central to memory formation and spatial navigation-is highly temperature sensitive ^7^. Prior work has shown that hippocampal physiology is sensitive to temperature changes ^7^. In freely moving rats, hippocampal temperature can rise by over 3 °C during behavior ^24^. In contrast, during prolonged electrical activation of hippocampal afferents in anesthetized rats, hippocampal temperature increases were modest (≤ ~0.6 °C). ^25^ Even moderate temperature shifts can alter electrophysiological properties: warmer temperatures speed axonal conduction and neurotransmitter release, while slightly reducing spike amplitude and modifying synaptic dynamics ^7,26^. Brain temperature also modulates quantitative features of hippocampal sharp-wave ripples, underscoring the sensitivity of hippocampal network dynamics to thermal state ^27^. Despite this recognized sensitivity, a major experimental gap remains: most thermometry approaches lack the temporal responsivity and spatial locality needed to resolve rapid, activity-locked temperature transients and to separate intrinsic thermogenesis from vascular cooling *in vivo*^*28*^.

Here, we directly measured temperature changes *ex vivo* within the apical dendritic region of hippocampal CA1 during presynaptically applied HFS using a micromachined thermocouple microprobe designed for rapid, localized thermometry^29^. Compared with conventional thermistors and macro-scale probes, micromachined sensors can achieve low thermal mass and short thermal time constants, enabling improved tracking of fast temperature dynamics that are otherwise temporally averaged. In this preparation, we isolate intrinsic activity-dependent heat generation from blood-flow-mediated heat removal, providing a direct measurement of circuit-driven thermogenesis during synchronized activation. By comparing responses in control conditions and during blockade of action potentials with tetrodotoxin, we aimed to quantify the magnitude of heating in CA1 due to synaptic signaling and to relate these thermal dynamics to established principles of neural energetics and brain temperature regulation.

## Materials and Methods

### Device Fabrication and Calibration

The micromachined thermocouple probes share the same overall architecture as previously reported and consist of a ~1.5 mm-long shank (~40 μm wide) ^29^. In the newer designs the shank tapers to a micrometer-scale distal tip (SEM of representative design shown in ***Fig. 1a***; tip width ≈6 μm over the final ~30 μm). To facilitate tissue penetration, the distal tip incorporates a triangular profile. In the design shown in ***Fig. 1a***. the thermocouple sensing junction is positioned near proximal to the tip and has an active junction area of approximately 3 × 4.5 μm. Micromachined fine microneedle thermocouple probes were fabricated on 4-inch silicon-on-insulator (SOI) wafers (10 μm device layer / 0.5 μm buried oxide (BOX) / 675 μm handle), following the process described in our previous work ^29^, with minor layout/dimensional variations across design iterations. Wafers were cleaned in NanoStrip (CMC Materials) at 60 °C for 10 min, followed by BF2 ion implantation (Coherent Corp., San Jose, CA) using conditions reported previously. Dopants were activated by furnace annealing at 850 °C for 30 min (Tempress), which reduced the sheet resistance from 9450 ± 250 to 291 ± 3 Ω/□. A 100 nm front-side silicon oxide layer was deposited by plasma enhanced chemical vapor deposition (PECVD) (Applied Materials P5000) to electrically isolate the doped silicon (***Fig. 1b: step 1***). Contact regions were patterned in ~2 μm SPR2000-3.0 photoresist (Kayaku Advanced Materials) using a stepper (GCA AS200 AutoStep) and transferred into the oxide by reactive oxide etching (LAM9400), followed by resist stripping and a NanoStrip (CMC Materials) clean. Metal features were then defined using a LOR10 B/S1813 (Kayaku Advanced Materials) liftoff stack patterned with the GCA AS200 AutoStep stepper (***Fig. 1b: step 2***). After a brief 1:20 buffered hydrofluoric acid dip (30 s) to remove native oxide, 40 nm chromium / 160 nm gold (Cr/Au) was deposited by evaporation (EvoVac, Angstrom Engineering) and lifted off in Remover PG to form the metallization (***Fig. 1b: step 2***). Thermocouple device geometries were subsequently patterned in SPR2000-3.0 using the GCA AS200 AutoStep stepper and transferred through the PECVD oxide (LAM9400) and into the 10 μm device silicon layer by deep reactive ion etching (DRIE) (STS Pegasus 4). Resist was stripped and wafers were cleaned in NanoStrip. A 30 nm aluminum oxide (Al_2_O_3_) insulation layer was deposited by atomic layer deposition (ALD) (Oxford OpAL ALD), and pad openings were defined in photoresist using the GCA AS200 AutoStep stepper and opened by buffered hydrofluoric acid etching (90 s), followed by resist stripping (***Fig. 1b: step 3***). For backside release, AZ12XT photoresist (MicroChemicals) was patterned on the backside using contact aligner (MA6BA6) and the wafer was temporarily bonded to a silicon carrier wafer. The handle silicon layer was etched by DRIE (STS Pegasus4) and the 0.5 μm BOX was opened using a plasma oxide etcher (STS APS DGDRIE). Finally, photoresist was stripped, the wafer was detached from the carrier, and the devices were cleaned in acetone and isopropanol (Transene) (***Fig. 1b: step 4***). The probe is a passive sensor during experiments (no current injection), minimizing self-heating artifacts.

**Fig. 1.**
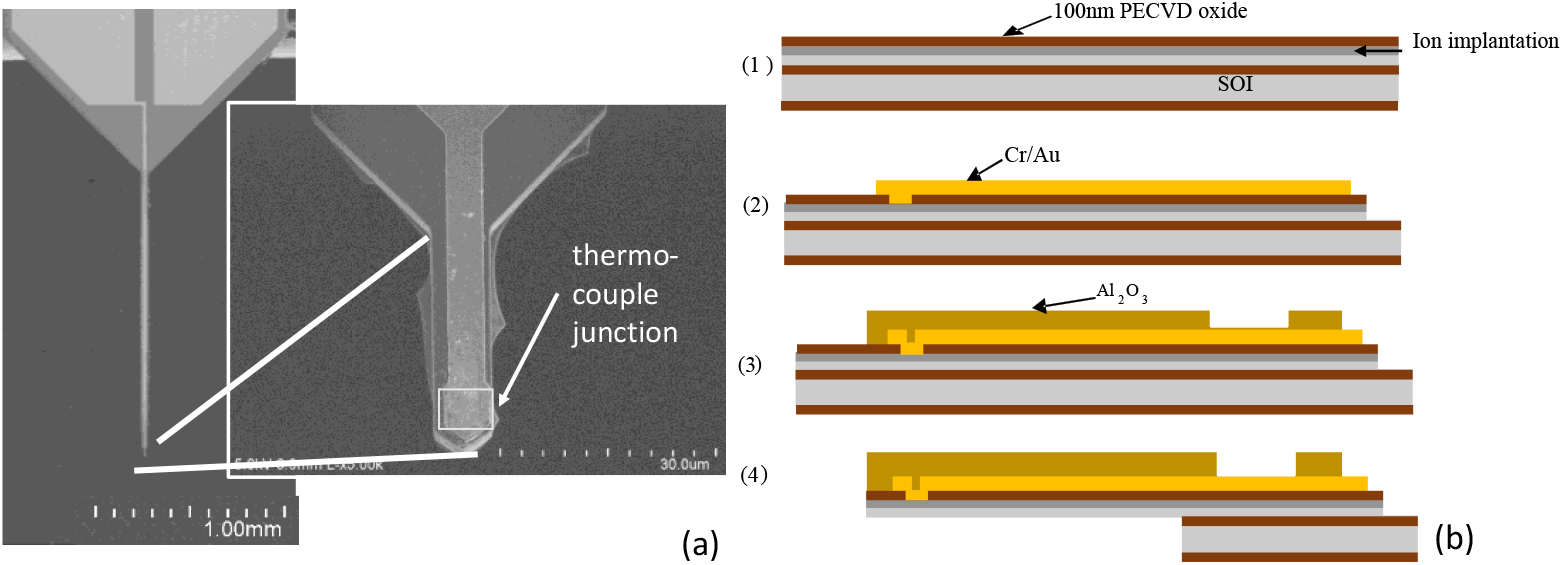
(a) Scanning electron microscope (SEM) images of the device; the inset shows a magnified view of the tapered tip region, which narrows to a 6 μm-wide section. The thermocouple junction area is ~ 3 × 4.5 μm. (b) Microfabrication process flow for the micromachined thermocouple.

Oil Droplet

The probes were calibrated *in vitro* in the set-up using a temperature controller (TC-344B, Warner Instruments, USA). During calibration, the controller incrementally increased the bath temperature up to +10 °C. The controller’s heating probe regulated the bath temperature, while an independent reference probe placed in the bath provided feedback measurements. To improve calibration accuracy and repeatability, we placed a small mineral oil droplet at the air-liquid interface around the probe entry point (***Fig. 2a***). This oil layer has a dual purpose: first, it suppresses capillary rise and meniscus “wicking” of aqueous solution along the probe shank, which can unintentionally bring portions of the probe near the reference junction/electrical interconnect into thermal contact with the heated bath and partially collapse the intended temperature difference between junctions; and second, it reduces evaporation at the interface, minimizing evaporative cooling that would otherwise bias the calibration curve. Together, these effects help ensure that the sensed thermovoltage reflects the intended junction temperature difference rather than artifacts from surface wicking or interface cooling. Calibration (***Fig. 2b***) produced a linear sensitivity of |S_eff| ≈ 292 μV/K (slope −292 μV/°C; R^2^ ≈ 0.996) for the Au-Si thermocouple. The negative slope arises from the chosen polarity/reference-junction convention (V_Au_ − V_Si_) with the reference junction near ambient, so increasing tip temperature reduces the measured ΔT relative to the reference and therefore reduces the output voltage.

**Fig. 2.**
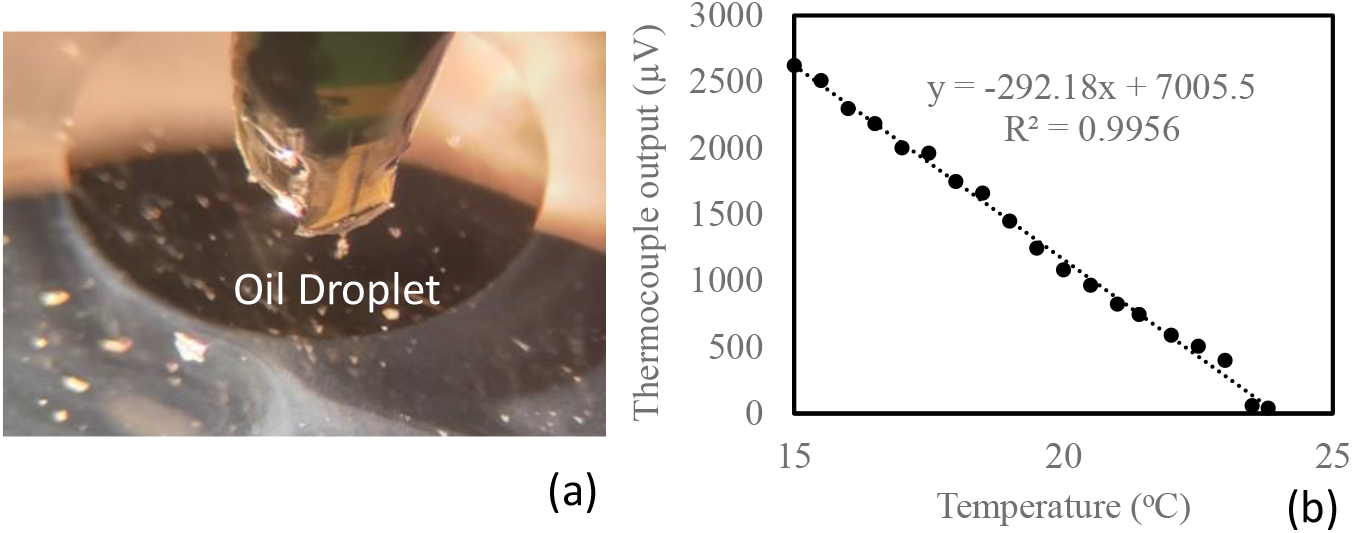
(a) Calibration setup using a thin mineral oil layer at the air-liquid interface to suppress meniscus formation/capillary rise and minimize evaporative cooling artifacts. (b) Thermocouple output vs temperature.

The thermal time constant of the probe in tissue was estimated using time-dependent finite-element simulations in COMSOL Multiphysics 5.5, following the modeling framework described in detail in our prior work^29^. Here, we repeated the same simulation workflow but updated the probe geometry to match the revised dimensions reported in this study (***Fig. 3a***). As in our previous analysis, simulations were used because the intrinsic sensor dynamics occur on timescales faster than the practical sampling limits of typical electrophysiology/data-acquisition hardware, consistent with common practice in micro-thermometry studies ^30,31^. Briefly, the model consisted of a 2 mm × 2 mm tissue block with the outer boundary held at ambient conditions (293 K) and a convective heat-flux boundary at the tissue-air interface using a heat-transfer coefficient of 10 W/(m^2^·K). The probe was initialized at 313 K and allowed to thermally equilibrate with the surrounding tissue. Material properties for silicon (Si), gold (Au), silicon dioxide (SiO_2_), aluminum oxide (Al_2_O_3_), and tissue were taken from COMSOL’s built-in libraries (***Fig. 3a***). Using these geometries, the estimated thermal time constant from ***Fig. 3b*** was ~8.5 μs for the updated probe layout and ~57 μs for the previously reported layout ^29^. Because both time constants are orders of magnitude shorter than a ~1 ms action potential, the measurement bandwidth is expected to be limited by tissue heat diffusion rather than probe response time.

**Fig. 3.**
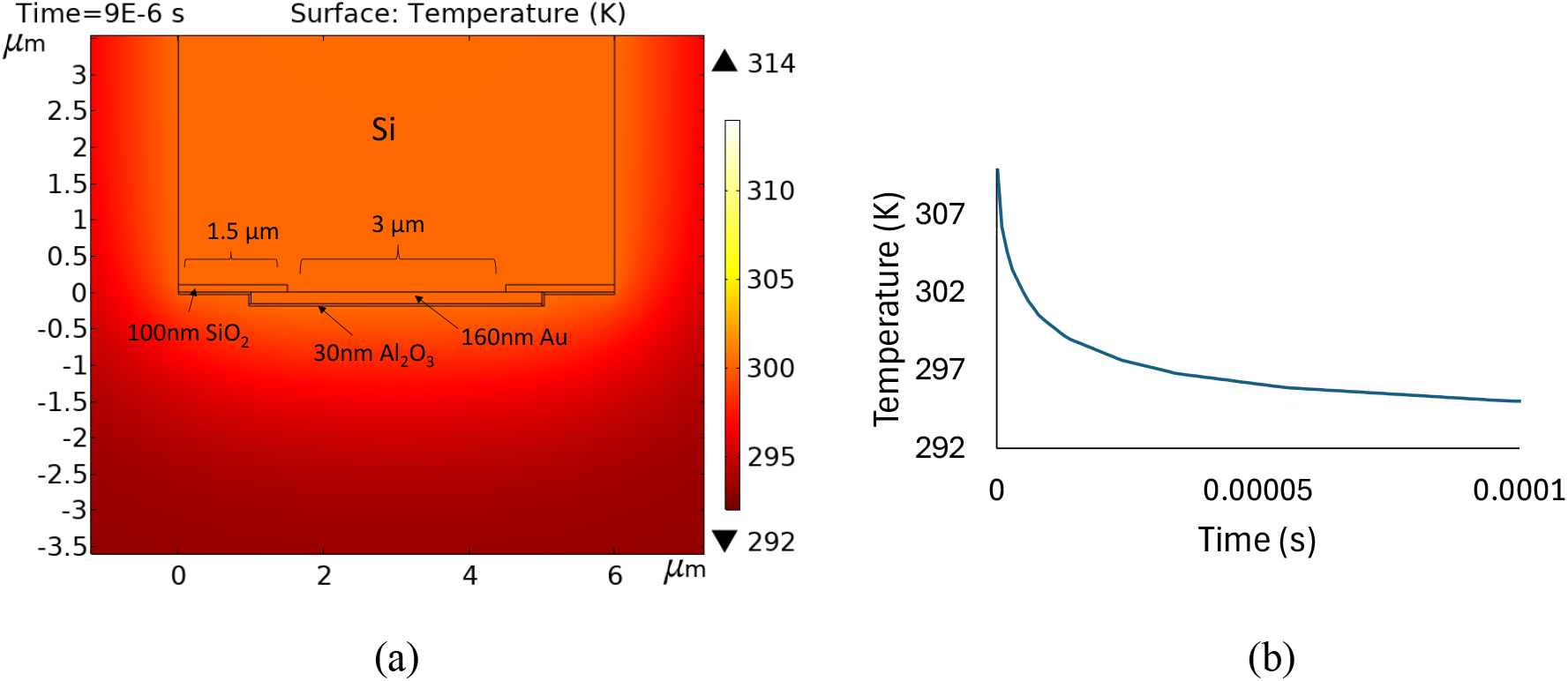
Finite-element thermal simulations. (a) Magnified view of the simulated temperature field near the probe tip at 9 10^−6^ s. (b) Simulated temperature transient at the Au-Si thermocouple junction, plotted as temperature versus time.

### Hippocampal Slice Preparation

4-8-week-old mice were deeply anesthetized with isoflurane and decapitated. The brain was rapidly removed and chilled in cutting artificial cerebrospinal fluid (ACSF) containing (in mM): N-methyl-D-glucamine 93, HCl 93, KCl 2.5, NaH_2_PO_4_ 1.2, NaHCO_3_ 30, HEPES 20, glucose 25, sodium ascorbate 5, thiourea 2, sodium pyruvate 3, MgSO_4_ 10, and CaCl_2_ 0.5, pH 7.4. The brain was embedded in 2% agarose and coronal slices (400 µm thick) were made using a Compresstome (Precisionary Instruments)^32^. Brain slices were allowed to recover at 31 ± 1 °C in cutting solution for 30 minutes and thereafter at room temperature in holding ACSF, containing (in mM): NaCl 92, KCl 2.5, NaH_2_PO_4_ 1.2, NaHCO_3_ 30, HEPES 20, glucose 25, sodium ascorbate 5, thiourea 2, sodium pyruvate 3, MgSO_4_ 2, and CaCl_2_ 2, pH 7.4. After at least 1 hour of recovery the slices were individually transferred to a submersion recording chamber and continuously perfused (2-3 ml/min) with ACSF containing (in mM): NaCl 124, KCl 2.5, NaH_2_PO_4_ 1.2, NaHCO_3_ 24, HEPES 5, glucose 12.5, MgSO_4_ 2, and CaCl_2_ 2, pH 7.4 at room temperature^33^. Slices were visualized under a stereomicroscope to target the CA1 region for electrode placement.

### Electrophysiological Stimulation and Recording

A concentric bipolar stimulating electrode (FHC, USA) was placed in the Schaffer collateral pathway (stratum radiatum near the CA3/CA1 border) to activate synaptic inputs to CA1. HFS protocols were used to evoke intense synaptic activity. The standard protocol consisted of a 1 second train at 100 Hz (100-μs pulse width, biphasic square pulses). This HFS train is comparable to those used to induce long-term potentiation and elicits robust multi-synaptic activation in CA1 ^34^. To monitor neuronal activity, a glass microelectrode (filled with ACSF, 2 MΩ resistance) was positioned in the CA1 pyramidal cell layer to record extracellular field potentials. Recordings were performed at room temperature using a Multiclamp 700A amplifier (Molecular Devices). Baseline stimulus-response properties were confirmed with test pulses (0.033 Hz) prior to high-frequency stimulation. In some experiments, we applied tetrodotoxin (TTX, 1 μM) to the bath for at least 10 minutes to block voltage-gated sodium channels and thus action potential firing^35^. This allowed us to stimulate the tissue under otherwise identical conditions but without neuronal spiking or synaptic transmission, isolating any non-physiological thermal artifact. Data acquisition (filtered at 10 kHz and digitized at 10 kHz) and analysis were performed with pClamp 11 software (Molecular Devices). Figure 4 shows the electrophysiology-thermometry setup and electrode/probe locations in the slice.

**Fig. 4.**
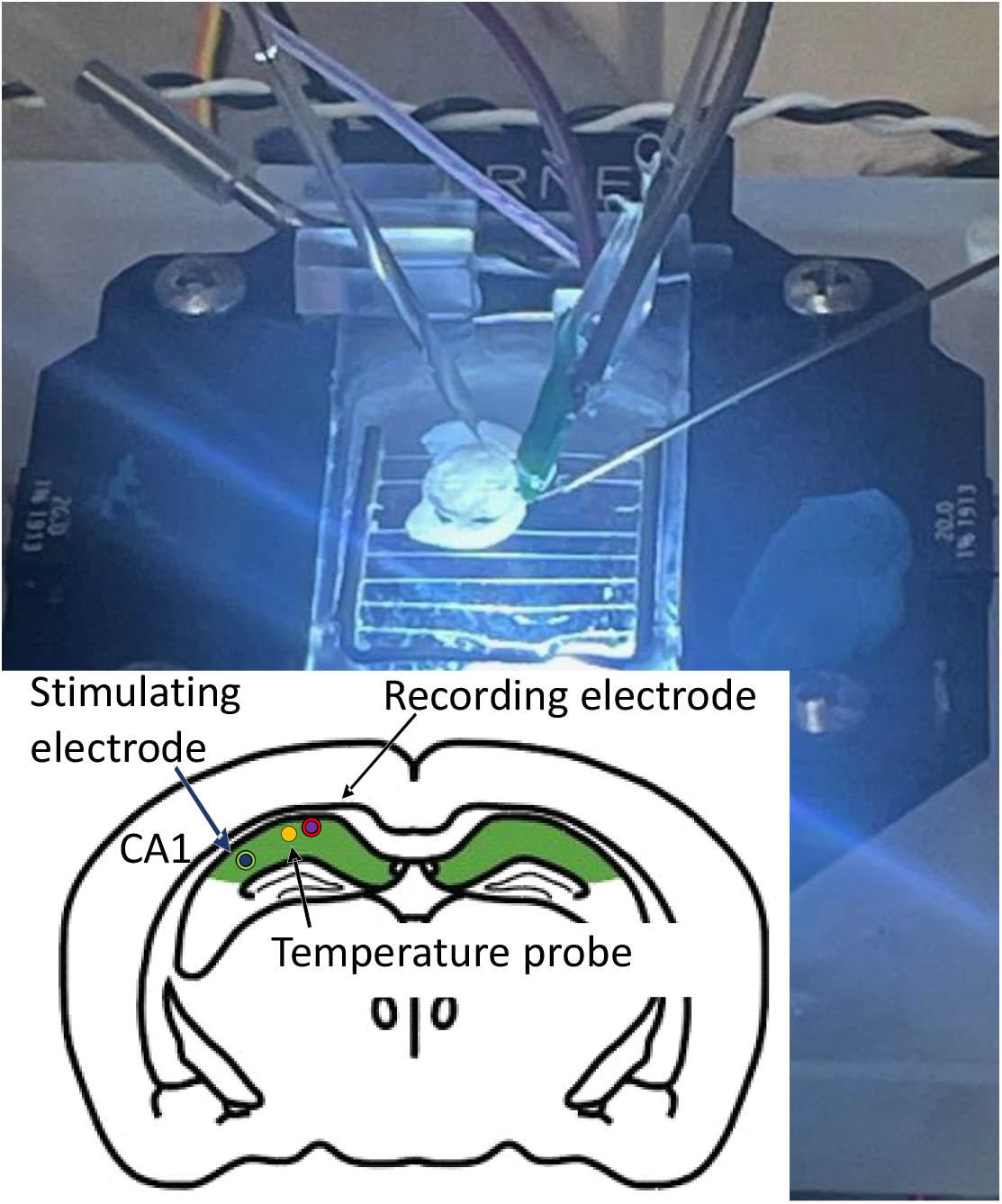
Electrophysiology-thermometry setup for hippocampal slice experiments. Top: Photograph of the recording chamber showing placement of the temperature probe in the slice together with the stimulation and recording electrodes. Bottom: Schematic indicating electrode locations relative to CA1.

### Temperature Measurements, Data Collection and Analysis

During each experiment, baseline temperature was recorded for at least 30 s prior to stimulation. The 100 Hz stimulation train was then delivered, and temperature recording continued for several minutes post-stimulation to capture the full return to baseline. Field potential recordings were synchronized to the temperature data to correlate electrical activity with thermal changes. Temperature traces were filtered to reduce high-frequency noise (using a 1 Hz low-pass digital filter) and analyzed to determine the peak change (ΔT) relative to the pre-stimulation baseline. Trials were repeated in each slice (with inter-train intervals >5 min to allow temperature to normalize). For TTX conditions, the stimulation protocol was repeated after drug perfusion. We quantified the mean peak ΔT for control vs. TTX conditions across slices. Statistical analysis was performed using a paired two-tailed Student’s t-test to assess the effect of TTX on the temperature response, with p < 0.05 considered significant. All data are reported as mean ± standard error of the mean (SEM).

## Results

### Baseline and Stimulus-Induced Temperature Changes

Prior to stimulation, slices maintained a stable baseline temperature (within ±0.02 °C), indicating minimal drift under our recording conditions. High-frequency synaptic stimulation of CA1 produced a rapid and significant increase in local tissue temperature. As shown in **Fig. 5a**, the 100 Hz, 1 s train elicited a temperature rise of approximately 0.6 °C in the apical dendritic layer of CA1 region (in this example, from 23.6 °C baseline to ~24.2 °C). The temperature began to increase shortly after the onset of the stimulus train and reached a peak of ΔT = 0.54 ± 0.07 °C (n = 4 slices) within a few seconds of train termination (***Fig. 5a***). This heat increase was transient; after the stimulus, the temperature returned toward baseline over the next few seconds. Figure 5a indicates the stimulation period, during which most of the temperature change occurred. Notably, the rise in temperature closely tracked the period of intense neural activity. We also recorded robust electrophysiological responses during these trials (black lines): the HFS train evoked a sustained burst of population spikes and field excitatory postsynaptic potentials (fEPSPs) in CA1, confirming that neurons were actively firing throughout the stimulation. The temporal profile of the temperature increase roughly corresponded to the duration of neural firing, with peak heat generation occurring at the end of the high-frequency burst. Consistent with recovery-phase thermogenesis, the thermal transient persisted beyond the stimulus train, decaying back to baseline on a slower timescale than the electrical activity itself.

**Fig. 5.**
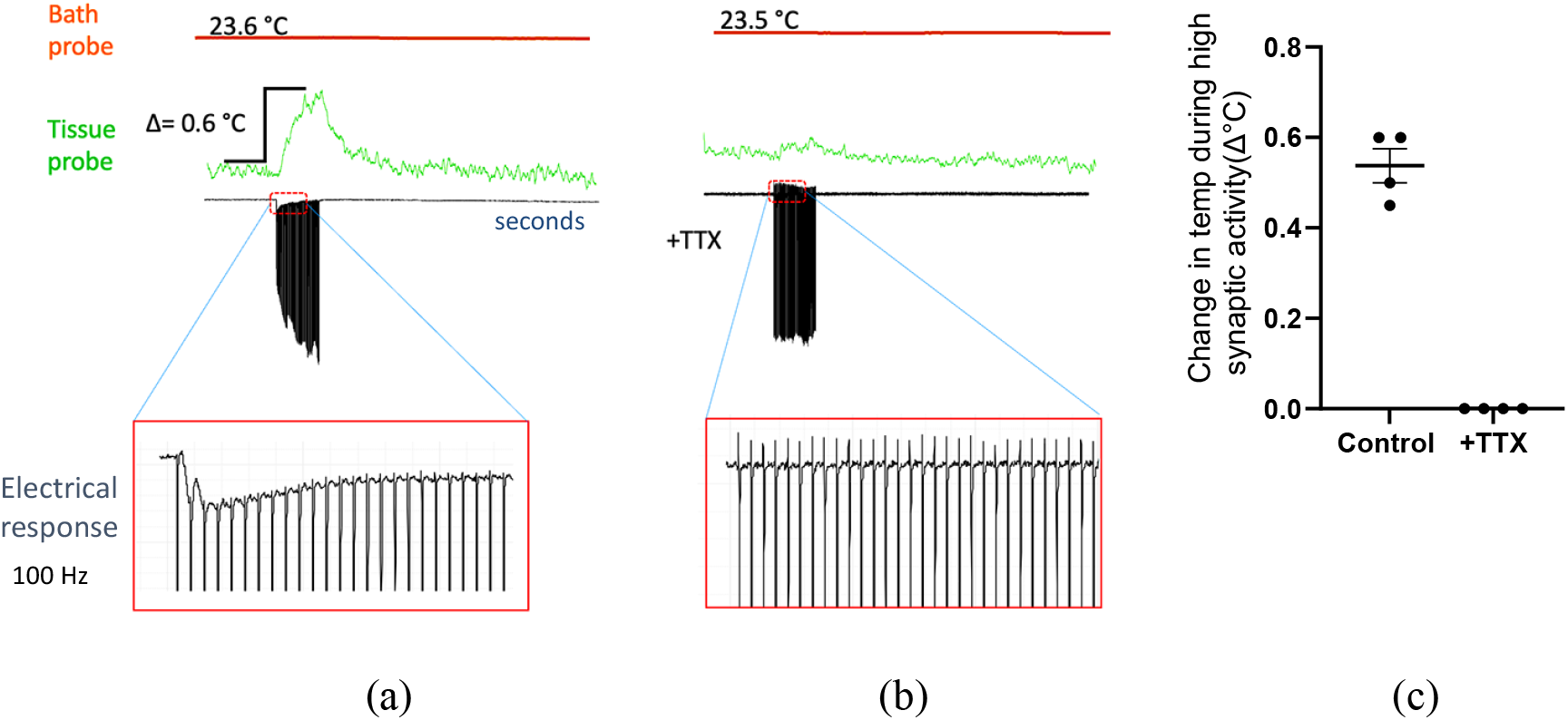
Local temperature responses in hippocampal CA1 during high-frequency stimulation. (a) In control slices, a 100 Hz stimulation (1 s) evoked a rapid rise in tissue temperature of about 0.6 °C. The green trace shows the temperature change (ΔT), with temperature peaking shortly after the stimulus train and gradually returning to baseline. (b) In the presence of 1 μM TTX to block neuronal firing, the same 100 Hz stimulation produced no significant temperature change (green trace) ^35^. Minor fluctuations remained within the noise level (~0.02 °C), indicating that the heat observed in (a) was due to action-potential-driven synaptic activity. (c) Four repetitions of stimulation with and without TTX.

In marked contrast, when neuronal activity was pharmacologically silenced by TTX: no meaningful temperature increase was detected. **Fig. 5b** shows the temperature trace during an identical 100 Hz stimulation in the presence of TTX. The temperature remained essentially flat, with only negligible noise-level variations. Field potential recordings confirmed that the HFS stimulus failed to elicit any neuronal activity in TTX-treated slices. Thus, the absence of a temperature change under TTX indicates that the heating observed in control conditions was intrinsically linked to neuronal firing and synaptic transmission. In other words, the tissue did not warm up simply from the delivery of electrical pulses or any Joule heating of the medium; it required active physiological metabolism. Comparing across conditions, the difference in peak ΔT between control and TTX was highly significant (p < 0.001). This provides strong evidence that the temperature rise in CA1 was a direct consequence of high-frequency neural activity. No other artifacts (such as external heating) were present in our *in vitro* setup that could account for this increase. We also verified that baseline slice temperature was stable over time (<0.05 °C drift over 10 min) and that repeated stimulations yielded consistent temperature responses after adequate recovery. Together, these results demonstrate localized thermogenesis in hippocampal tissue during periods of intense synaptic activation, and confirm that it depends on action potential firing and synaptic function.

## Discussion

Our findings provide direct experimental evidence that high-frequency synaptic activity can induce measurable local thermogenesis in hippocampal tissue. In a controlled slice preparation, a brief 100 Hz stimulus train elicited a transient ~0.54 °C rise in temperature within the CA1 region, whereas blocking neuronal activity with TTX abolished this temperature increase. These results establish a causal link between neural firing and heat production at the tissue level^1,13^. The magnitude of the observed heating, while modest, is remarkable given the small volume of tissue involved and the short stimulation duration. It underscores that even a second of intense spiking activity by hippocampal neurons is sufficient to measurably warm the surrounding brain tissue.

This localized temperature elevation is attributable to the surge in metabolic activity accompanying synaptic transmission and action potential generation. Neurons rapidly consume ATP to fuel ion pumps (Na^+^/K^+^-ATPases) that restore ionic gradients disrupted during spiking. The hydrolysis of ATP releases energy, a portion of which dissipates as heat^1,14,15^. Enzymatic processes and mitochondrial oxidative metabolism also contribute to heat production during heightened activity ^1^. Our TTX control confirms that the heating requires active metabolism: without ion fluxes and synaptic currents, no significant temperature change occurs. Thus, the hippocampus behaves like a heat engine during high-frequency firing, converting chemical energy (glucose/ATP) into thermal energy.

Beyond neuronal ion pumping, the observed thermogenesis likely includes a significant contribution from glial metabolism. In the hippocampus, high-frequency stimulation triggers massive glutamate release, the clearance of which by astrocytes is an energy-intensive process requiring the hydrolysis of ATP to transport glutamate against its concentration gradient^17,18^. The astrocytic glutamate-glutamine cycle and associated glucose uptake provide a secondary, prolonged heat source that contributes to the ‘recovery’ phase of the temperature transient^19,36^. This metabolic cooperation suggests that the recorded temperature rise is a network-level signature of coupled neuronal and glial activity. This interpretation is consistent with the biphasic heat framework: the measured temperature transient at tissue scale is expected to weight recovery metabolism more strongly than the instantaneous “initial heat” component.

These data align with observations from in vivo studies of brain temperature. The ~0.5-0.6 °C increase we report in an isolated hippocampal slice is comparable to the local temperature rises recorded in intact brains during intense neural activation. For instance, direct thermistor measurements in rats have shown that stimulating hippocampal afferents (perforant path) causes gradual temperature increases on the order of 0.1-0.5 °C in the dentate gyrus ^24,25^. In awake, behaving animals, natural stimuli and behavioral arousal can drive brain temperature elevations of similar or greater magnitude^1,13^. Even mild sensory events cause small but reliable temperature upticks in corresponding brain regions ^4,12^. Notably, brain temperature changes often precede and exceed any concomitant changes in core body temperature^1,13^, indicating their origin is neural rather than systemic. Our demonstration that eliminating neural firing (with TTX) prevents the heat rise reinforces the interpretation that neural activity is the primary driver of these temperature fluctuations.

There are, however, important differences between our *in vitro* slice preparation and the intact brain environment. *In vivo*, cerebral blood flow plays a critical role in dissipating heat. The bloodstream effectively removes heat from active brain regions, analogous to a radiator cooling a car engine ^1,21 23,37^. In a slice, lacking blood perfusion, heat generated by neurons must diffuse through the tissue and into the perfusate. This could lead to slightly higher local temperature accumulation in slices for a given level of activity, or conversely, limit heat production if oxygen/substrate delivery becomes limiting. Our measured 0.54 °C rise likely represents a scenario of localized heating without vascular heat removal (***Fig. 5c***). Despite these differences, the consistency in scale between our results and *in vivo* data suggests that hippocampal thermogenesis is a robust phenomenon observable across conditions.

Mechanistically, the ability of neurons to generate heat is a natural consequence of their low thermodynamic efficiency. A substantial portion of the energy consumed during intense firing is not converted into electrical work, but lost as heat ^14,15^. Prior studies have pointed out that during high-frequency spiking, the energetic cost (in ATP) of maintaining ion gradients increases, which would proportionally increase heat output^1^. Additionally, mitochondrial respiration may become elevated to replenish ATP, and since oxidative phosphorylation is exothermic, it adds to the thermal load^1^. Our experimental design does not allow us to pinpoint the exact sources of the heat (ion pumps vs. other metabolism), but the requirement for spiking suggests a dominant contribution from processes linked to action potentials and synaptic currents. Interestingly, we noted that the temperature peaked just after the 1 s stimulus train and decayed over several seconds, mirroring the time course of post-tetanic metabolic activity (e.g., sustained Na^+^/K^+^-ATPase activity and recovery processes) rather than the instantaneous electrical activity. This lag and relaxation are consistent with the idea that heat continues to be produced as neurons restore homeostasis following intense firing, and then dissipates into the environment.

From a functional perspective, the local temperature transients we observed could have several implications for hippocampal physiology. Although a ~0.54 °C change is relatively small, prior work has shown that even minor temperature shifts can modulate neuronal and synaptic behavior ^7,26,27^. Warming of hippocampal tissue tends to increase membrane conductance and neurotransmitter release probability, leading to faster synaptic potentials and reduced synaptic delay ^7^. It also shortens action potential duration and decreases spike amplitude due to altered ion channel kinetics ^26^. In our experiments, the brief heating during the HFS might transiently enhance synaptic transmission or alter the excitability of CA1 neurons, potentially contributing to plasticity phenomena.

Our results also prompt considerations for extreme conditions. In normal physiological settings, the brain’s thermoregulatory mechanisms (like increased blood flow to active areas) likely prevent minor activity-related heat from accumulating pathologically. But if those mechanisms are overwhelmed or impaired, activity-driven thermogenesis could compound to dangerous levels. A striking example is the effect of psychostimulant drugs: these substances increase neural firing and can also reduce heat dissipation through peripheral vasoconstriction, potentially allowing for aberrantly rapid and pronounced increases in brain temperature. Amphetamine-like drugs can drive brain temperatures above 40 °C, contributing to blood-brain barrier disruption and cellular injury ^38,39^. Such drug-induced hyperthermia underscores how local heat production by neurons, when unchecked, can escalate to pathological outcomes. While our hippocampal slices were in a well-controlled environment, the fundamental process we observedneurons converting increased activity into heat-is the same process that can contribute to hyperthermia and potential brain damage in severe settings. This connection highlights the clinical relevance of understanding neural thermogenesis and suggests that brain temperature could serve as a readout of neural metabolic stress in contexts such as seizures or stimulant exposure.

There are several technical considerations and limitations in this study. First, our measurement approach provides a single-point average temperature in the tissue near the thermocouple tip. It does not resolve microscale temperature gradients that might exist in either the pre- or postsynaptic neuron, nor across the dendritic arbor. Information on the latter issue would be of particular interest, since the spread of dendritic heating some distance from activated synapses could have repercussions for synaptic associativity and synaptic capture. Advanced temperature-imaging approaches (including fluorescent nanothermometers) suggest that intracellular microdomains can exhibit larger transient heating during intense metabolic bursts^1,28^. Our results do not allow us to detect such local hotspots, and instead reflect the composite heating of many cells in the probe’s vicinity. Second, the use of an *in vitro* slice means we isolated the hippocampus from its normal environment. While this offers the advantages of accessibility and precise positioning of thermal probe and electrodes, it also removes factors like neurovascular coupling that *in vivo* would modulate both activity and heat dispersal. Future experiments could extend these findings by measuring temperature *in vivo* in the hippocampus during similar stimulation paradigms or by using imaging methods to map the spatial distribution of heat in slices. Another extension would be to examine prolonged or repetitive stimulation to determine whether small temperature increments can accumulate or trigger homeostatic responses.

In conclusion, this study demonstrates that localized thermogenesis accompanies high-frequency synaptic activity in the hippocampus. The CA1 region of hippocampal slices experienced reproducible temperature rises when its neuronal circuits were driven at 100 Hz, and this effect was strictly activity-dependent. These findings reinforce the view that the brain’s functional activity and thermal state are closely coupled^1^. Monitoring and controlling for such temperature changes is important in neurophysiological experiments because temperature can affect synaptic and circuit behavior ^7,26^. Beyond experimental considerations, elucidating how the brain dissipates or leverages activity-induced heat could shed light on neurovascular regulation and inform treatment strategies for conditions where brain hyperthermia is a risk. The interplay between neural activation, energy consumption, and heat production is a fundamental aspect of brain physiology that warrants further exploration.

## Acknowledgements

This work was supported by the National Science Foundation under award numbers 226930 and 2442758, the content is solely the responsibility of the authors and does not necessarily represent the official views of the National Science Foundation. The thermocouple probe was fabricated at the Lurie Nanofabrication Facilities at the University of Michigan. Hippocampal slice experiments were conducted at the Electrophysiology Core at the Icahn School of Medicine at Mount Sinai, New York.

## Author Contributions

V.P.: conceived/designed/executed/acquired/analyzed/retained hippocampal slice experiments and data, setup, preparation, in situ calibration, visualization, data curation. R.D.B.: conceived/designed slice experiments, contributed to slice experimental design/interpretation; oversight/resources. O.S.: probe/device design, device fabrication. I.C.T.: liquid calibration/validation. A.G.: conceived/designed experiments; co-designed probe; liquid calibration/validation; wrote original draft; supervision. All authors: review and revision.

## References

1. Kiyatkin EA. Brain temperature and its role in physiology and pathophysiology: Lessons from 20 years of thermorecording. Temperature 2019;6(4):271–333.

2. Abrams R, Hammel H. Hypothalamic temperature in unanesthetized albino rats during feeding and sleeping. American Journal of Physiology-Legacy Content 1964;206(3):641–646.

3. Delgado J, Hanai T. Intracerebral temperatures in free-moving cats. American Journal of Physiology-Legacy Content 1966;211(3):755–769.

4. Baker M, Frye FM, Millet V. Origin of temperature changes evoked in the brain by sensory stimulation. Experimental Neurology 1973;38(3):502–519.

5. Kiyatkin EA. Brain temperature: from physiology and pharmacology to neuropathology. Handbook of clinical neurology 2018;157:483–504.

6. Wang H, Wang B, Normoyle KP, et al. Brain temperature and its fundamental properties: a review for clinical neuroscientists. Frontiers in neuroscience 2014;8:307.

7. Andersen P, Moser EI. Brain temperature and hippocampal function. Hippocampus 1995;5(6):491–498.

8. Serota H, Gerard R. Localized thermal changes in the cat’s brain. Journal of neurophysiology 1938;1(2):115–124.

9. Mellergard P, Nordström C-H. Epidural temperature and possible intracerebral temperature gradients in man. British journal of neurosurgery 1990;4(1):31–38.

10. Kovalzon V. Brain temperature variations during natural sleep and arousal in white rats. Physiology & Behavior 1973;10(4):667–670.

11. Smirnov MS, Kiyatkin EA. Fluctuations in central and peripheral temperatures associated with feeding behavior in rats. American Journal of Physiology-Regulatory, Integrative and Comparative Physiology 2008;295(5):R1415–R1424.

12. McElligott J, Melzack R. Localized thermal changes evoked in the brain by visual and auditory stimulation. Experimental neurology 1967;17(3):293–312.

13. Kiyatkin EA, Brown PL, Wise RA. Brain temperature fluctuation: a reflection of functional neural activation. European Journal of Neuroscience 2002;16(1):164–168.

14. Ritchie JM. Energetic aspects of nerve conduction: the relationships between heat production, electrical activity and metabolism. Progress in biophysics and molecular biology 1973;26:147–187.

15. Laughlin SB, de Ruyter van Steveninck RR, Anderson JC. The metabolic cost of neural information. Nature neuroscience 1998;1(1):36–41.

16. Abbott BC, Hill AV, Howarth J. The positive and negative heat production associated with a nerve impulse. Proceedings of the Royal Society of London Series B-Biological Sciences 1958;148(931):149–187.

17. Danbolt NC. Glutamate uptake. Progress in neurobiology 2001;65(1):1–105.

18. Pellerin L, Magistretti PJ. Glutamate uptake into astrocytes stimulates aerobic glycolysis: a mechanism coupling neuronal activity to glucose utilization. Proceedings of the National Academy of Sciences 1994;91(22):10625–10629.

19. Attwell D, Laughlin SB. An energy budget for signaling in the grey matter of the brain. Journal of Cerebral Blood Flow & Metabolism 2001;21(10):1133–1145.

20. Howarth C, Gleeson P, Attwell D. Updated energy budgets for neural computation in the neocortex and cerebellum. Journal of Cerebral Blood Flow & Metabolism 2012;32(7):1222–1232.

21. Hayward JN, Baker MA. Role of cerebral arterial blood in the regulation of brain temperature in the monkey. American Journal of Physiology-Legacy Content 1968;215(2):389–403.

22. Pennes HH. Analysis of tissue and arterial blood temperatures in the resting human forearm. Journal of applied physiology 1948;1(2):93–122.

23. Sukstanskii AL, Yablonskiy DA. Theoretical model of temperature regulation in the brain during changes in functional activity. Proceedings of the national academy of sciences 2006;103(32):12144–12149.

24. Moser E, Mathiesen l, Andersen P. Association between brain temperature and dentate field potentials in exploring and swimming rats. Science 1993;259(5099):1324–1326.

25. Moser EI ML. Relationship between neuronal activity and brain temperature in rats. NeuroReport 1996;7(11):1876.

26. Thompson SM, Masukawa LM, Prince DA. Temperature dependence of intrinsic membrane properties and synaptic potentials in hippocampal CA1 neurons in vitro. The Journal of neuroscience 1985;5(3):817.

27. Petersen PC, Vöröslakos M, Buzsáki G. Brain temperature affects quantitative features of hippocampal sharp wave ripples. Journal of Neurophysiology 2022;127(5):1417–1425.

28. Zhu H, Xu H, Zhang Y, et al. Exploring the Frontiers of Cell Temperature Measurement and Thermogenesis. Advanced Science 2025;12(1):2402135.

29. Srivannavit O, Joshi R, Zhu W, et al. Design, fabrication, and calibration of a micromachined thermocouple for biological applications in temperature monitoring. Biosensors and Bioelectronics 2025;267:116835.

30. Wang C, Xu R, Tian W, et al. Determining intracellular temperature at single-cell level by a novel thermocouple method. Cell research 2011;21(10):1517.

31. Rajagopal MC, Valavala KV, Gelda D, Ma J, Sinha S. Fabrication and characterization of thermocouple probe for use in intracellular thermometry. Sensors and Actuators A: Physical 2018;272:253–258.

32. Serafini RA, Farzinpour Z, Patel V, et al. Nucleus accumbens myocyte enhancer factor 2C mediates the maintenance of peripheral nerve injury–induced physiological and behavioral maladaptations. Pain 2024;165(12):2733–2748.

33. Godino A, Salery M, Minier-Toribio AM, et al. Dopaminoceptive D1 and D2 neurons in ventral hippocampus arbitrate approach and avoidance in anxiety. bioRxiv 2023.

34. Bliss TV, Lømo T. Long-lasting potentiation of synaptic transmission in the dentate area of the anaesthetized rabbit following stimulation of the perforant path. The Journal of physiology 1973;232(2):331–356.

35. Narahashi T, Moore JW, Scott WR. Tetrodotoxin blockage of sodium conductance increase in lobster giant axons. The Journal of general physiology 1964;47(5):965–974.

36. Sibson NR, Dhankhar A, Mason GF, Rothman DL, Behar KL, Shulman RG. Stoichiometric coupling of brain glucose metabolism and glutamatergic neuronal activity. Proceedings of the National Academy of Sciences 1998;95(1):316–321.

37. Yablonskiy DA, Ackerman JJ, Raichle ME. Coupling between changes in human brain temperature and oxidative metabolism during prolonged visual stimulation. Proceedings of the National Academy of Sciences 2000;97(13):7603–7608.

38. Kiyatkin EA. The hidden side of drug action: brain temperature changes induced by neuroactive drugs. Psychopharmacology 2013;225(4):765–780.

39. Kiyatkin EA. State-dependent and environmental modulation of brain hyperthermic effects of psychoactive drugs of abuse. Temperature 2014;1(3):201–213.

